# Evaluation of antimalarial properties from Azores deep-sea invertebrate extracts: a first contribution

**DOI:** 10.1101/030007

**Authors:** Silvia Lino, Marta Machado, Dinora Lopes, Vírgilio do Rosário, Ricardo S. Santos, Ana Colaço

## Abstract

Several deep-sea marine invertebrates were collected from hydrothermal vents, seamounts and cold coral assemblages in the North Atlantic Ocean, near the Azores islands. The effect of their lipid crude extracts against two strains of malaria parasite *P. falciparum* (Dd2 and 3D7) was measured *in vitro* in order to establish the potential of these invertebrates as new sources for antimalarial compounds. Extracts that presented higher antimalarial activity potential were gorgonian *Callogorgia verticillata* and the hydrothermal vent shrimp *Mirocaris fortunata*, presenting the lowest value of IC50 and the highest selectivity index for both evaluated stains. To our knowledge this is the first report on the antimalarial activity of crude lipid extracts considering species collected at depths higher then 100 m.

## 1. Introduction

Only in Africa, malaria kills about two million every year, and is the leading cause of illness and death among children and pregnant women in many Sub-Saharan African countries (1). Several studies about therapeutic response showed that with the rise of chloroquine resistance, chloroquine efficacy fallen from nearly 100% to less than 25% in some endemic areas, resulting in an increase morbidity and mortality caused by malaria infection (1). Chloroquine (CQ) resistance in *Plasmodium falciparum* was first noticed in the late 1950s in non-immune migrant workers in hyperendemic areas along the Thai–Cambodian border, and was followed by resistance in at least three further geographically separate regions: two in Africa and one in Brazil (2). This marked the beginning of an era of rapid evolution from mono- to multi-drug resistant parasites and within 20 years CQ resistance has spread to most of the world (2). By now it is of common understanding that drug resistance in parasites establishes rapidly, disseminates widely, it is a persistent global health threat and represents the major problem for the success of the malaria control programs, justifying the urgency in developing new drugs. To resolve the emergency of finding new alternatives for the malaria therapeutic it is of great interest that the search of the antimalarial properties focuses especially on the erythrocytic stages of the parasite life cycle.

New compounds, from unexpected sources can hold the key to fight against this resistance and several studies on marine lipid compounds as antimalarial agents have already shown promising results (3, 4, 5, 6). Tasdemir and collaborators (7) showed that lipid crude extracts of sponge *Agelas oroides* were good inhibitors (IC50 of 0.35 μg/ml) for *P. falciparum* FA synthase, an important enzyme for the parasite fatty acids production (source of energy and *building blocks* to integrate in cell membranes). This finding was particularly interesting since this particular type II FA (FASII) system is absent in humans, presenting thus low cytotoxicity on mammalian cells (4), one of the most important pharmaceutical profile to search in a drug formulation.

Nevertheless, the research on marine bioactive natural products, when compared to plant natural products is still in its infancy. Only in December of 2004, the first marine-derived compound was approved in the United States: the Ziconotide (Prialt; Elan Pharmaceuticals), a peptide originally discovered in a tropical cone snail, used for the treatment of pain (8). One of the biggest challenges is the research itself, as the marine animals are not as easily accessed as the terrestrial animals and plants. Specially deep-sea animals which requires the use of submersibles and/or remotely operated vehicles (ROVs), faces even greater difficulties as only a few scientific institutions worldwide have the capacity to access those depths (9). To our knowledge this is the first study to evaluate the antimalarial activity of deep-sea marine invertebrates collected in the Atlantic. Azores is a region in the northeast near the Mid-Atlantic Ridge, particularly interesting as it hosts a great diversity of deep-sea environments, with numerous hydrothermal vents, seamount ecosystems, sponge aggregations and cold water coral assemblages.

## 2. Results and Discussion

Total lipids were extracted from eight invertebrate species (Figure 1) collected in North Atlantic Ocean near Azores islands: mussel *Bathymodiolus azoricus*, bristle worm *Branchipolynoe seepensis* and shrimp *Mirocaris fortunata* from hydrothermal vents communities; sponges *Neophrissospongia nolitangere, Petrosia* sp. and *Leiodermatium* cf. *pfeifferae;* and cold water gorgonian corals *Callogorgia verticillata* and *Dentomuricea* sp. (both common components of deep-sea communities in the Azores waters). From these animals, eleven crude extracts for total lipids were obtained, with quantities ranging from 56 to 319 mg. All extracts were standardized to have an initial concentration of 10 mg/ml in DMSO (dimethyl sulfoxide) as stock solutions for antimalarial essays. To ensure total extract dissolution in DMSO, a sonication bath with warm water was used previously to preparation of concentrations series with dilutions in RPMI medium.

**Figure 1.**
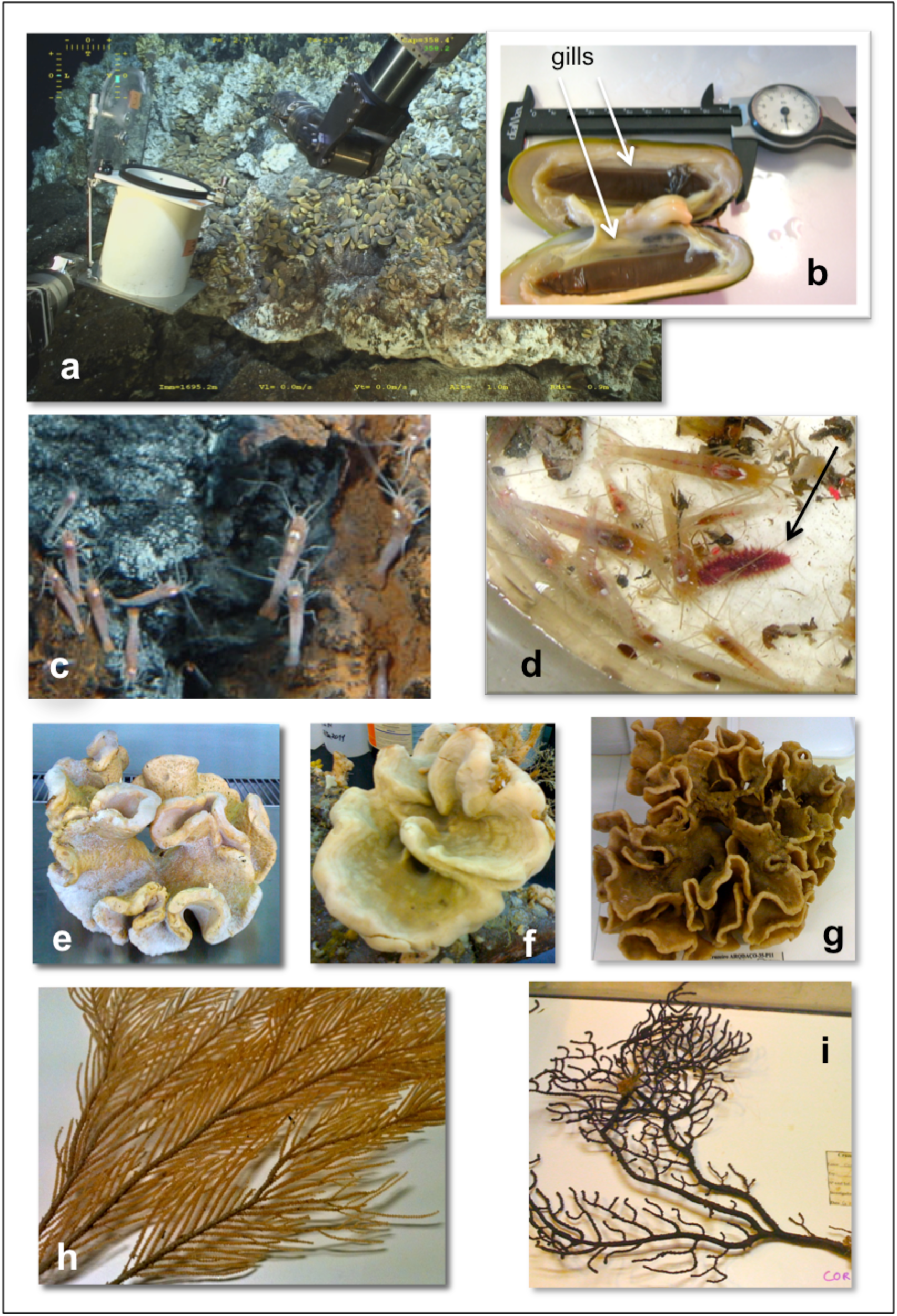
Biological samples used in this study **(a)** Mussel *B. azoricus* collection at Lucky Strike hydrothermal vent (copyright: Victor 6000, IFREMER) and **(b)** opened mussel *B. azoricus* with gills (indicated by white arrows); **(c)** Hydrothermal vent shrimp *M. fortunata* (copyright: Victor 6000, IFREMER); **(d)** red polychaete *B. seepensi* (indicated by black arrow) **(e)** sponge *N. nolitangere*; **(f)** sponge *Petrosia sp*; **(g)** sponge *L.* cf. *pfeifferae*; **(h)** gorgonian *C. verticillata*; and **(i)** gorgonian *Dentomuricea* sp.

Hydrothermal vent mussels are known for harboring chemosynthetic symbiotic bacteria that allows them to live in those environments (10). To test the hypothesis if tissues harboring symbiotic bacteria would present different results in terms of *P. falciparum* inhibition than tissues without bacteria, all mussels were also dissected in gills (tissues that contain symbiotic bacteria), muscle and digestive glands and extracted individually. From the pharmacologic point of view, the parameter considered to report the activity in terms of parasite inhibition is the concentration of an inhibitory agonist that reduces a response by 50% of maximal inhibition that can be attained, the IC50 (11). Antimalarial activity was evaluated using the WHO microtest methodology and IC50s were determined using GraphPad Prism 5 software (Table 1).

**Table 1.** Sample information, cytotoxicity and selectivity index for all crude extracts investigated. Evaluation of antimalarial activity is expressed in terms of IC50 (μg/ml). Selectivity index (SI) is defined as ratio of cytotoxicity (IC50) to antiplasmodial activity (IC50)

| Invertebrate/tissue<br>(specie) | Collection: |  | IC<sub>50</sub> (µg/ml) ± SD |  |  | Selectivity Index |  |
| --- | --- | --- | --- | --- | --- | --- | --- |
|  | Site | Depth<br>(m) | *P. falciparum*<br>strains |  | HepG2<br>cells | HepG2/<br>3D7 | HepG2/<br>Dd2 |
|  |  |  | 3D7 | Dd2 |  |  |  |
| Mussel gills<br>(*B. azoricus*) | *Lucky Strike*<br>hydrothermal<br>vent | 1700 | 59.7±<br>3.28 | 71.8±<br>1.64 | 259.3±<br>9.63 | 4.34 | 3.61 |
| Mussel digestive<br>glands<br>(*B. azoricus*) | *Lucky Strike*<br>hydrothermal<br>vent | 1700 | 100.4±<br>4.97 | 107.9±<br>6.70 | 95.3±<br>10.80 | 0.95 | 0.88 |
| Mussel muscle<br>(*B. azoricus*) | *Lucky Strike*<br>hydrothermal<br>vent | 1700 | 158.0±<br>11.03 | 155.6±<br>2.40 | 331.3±<br>19.65 | 2.10 | 2.13 |
| Polychaete<br>(*B. seepensis*) | *Lucky Strike*<br>hydrothermal<br>vent | 1700 | 87.0±<br>19.97 | 91.7±<br>17.97 | 266.2±<br>27.51 | 3.06 | 2.90 |
| Shrimp<br>(*M. fortunata*) | *Lucky Strike*<br>hydrothermal<br>vent | 1700 | 76.6±<br>6.33 | 78.3±<br>1.79 | 515.1±<br>24.61 | 6.72 | 6.58 |
| Sponge<br>(*N. nolitangere*) | Terceira<br>Island slope | 102 | 83.2±<br>7.65 | 66.9 ±<br>3.94 | 304.8±<br>11.87 | 3.66 | 4.56 |
| Sponge 1<br>(*Petrosia* sp.) | Faial Island<br>slope | 216 | 115.8±<br>6.71 | 100.5±<br>0.96 | 56.7±<br>7.66 | 0.49 | 0.56 |
| Sponge 2<br>(*Petrosia* sp.) | Princesa<br>Alice<br>seamount | 300 | 116.5±<br>17.11 | 105.0±<br>10.88 | 483.0±<br>22.30 | 4.15 | 4.60 |
| Sponge<br>(*L. cf. pfeifferae*) | Açor<br>seamount | 384 | 56.6±<br>3.79 | 62.4±<br>0.40 | 107.6±<br>8.35 | 1.90 | 1.72 |
| Gorgonia coral<br>(*C. verticillata*) | Açor<br>seamount | 409 | 44.0±<br>6.10 | 46.6±<br>7.64 | 268.6±<br>35.6 | 6.10 | 5.77 |
| Gorgonia coral<br>(*Dentomuricea* sp.) | Condor<br>seamount | 193 | 60.1±<br>12.80 | 59.3±<br>5.00 | 281.8±<br>17.61 | 4.69 | 4.75 |

In this study, crude extracts that were considered the most promising were the ones with both low IC50 for the antiplasmodial activity, high IC50 for the citotoxity essay and high values for the selectivity index. The hydrothermal shrimp and gorgonia *Callogorgia verticillata* were the crude extracts that fulfilled these requirements for both strains, with selectivity index of 6.72 and 6.10 for strain 3D7 and 6.58 and 5.77 for dD2, respectively (see Table 1).

Surprisingly, the most active extracts were not the sponges, already described in other studies as having antimalarial compounds (3, 7) but an animal with hydrothermal origin and a deep-sea gorgonia coral. To our knowledge this is the first report on the antimalarial activity from deep-sea hydrothermal vent species. The vent mussel *B. azoricus*, that on previous studies presented antibacterial activity (12) and anticancer activity (13) does not show as promising antimalarial activity as others tested animals. However, the gills from the mussel, that host 2 types of symbiotic bacteria (14) presents a fair low IC50 value (59.7 and 71.8 μg/ml). This indicates that the origin for the antiplamodial activity exhibited by *B. azoricus* might be from the symbionts and not exactly from the mussel tissue (with IC50 158 μg/ml for 3D7 and 156 μg/ml for Dd2). Considering the antimalarial potential of hydrothermal vent animals, the mussel gill presents a lower value of SI then the shrimp (although IC50 is lower).

The IC50 calculated for lipid extracts evaluated in this study range from 44 to 158 μg/ml. Tasdemir and collaborators (7) refers to IC50 from crude extracts of *Agelas oroides* sponge against *P. falciparum* (*in vitro* essays) ranging from 3.9 to less then 50 μg/ ml. Although with high IC50 values, the tested extracts are lipid crude extract and not isolated compounds that usually refer to having lower IC50 values (7).

This work intends to provide the first insights on antimalarial potentials for deep-sea invertebrate lipid extracts. Lipids are known as bringing advantages in terms of pharmaceutical drug formulation, as these compounds are highly permeable in cells, a desirable ‘drug-like’ property for a compound (15). Next steps will be purification and isolation of active compounds in most promising extracts and *in vivo* activity assessment for compounds in order to draw conclusions on the potential usages of these marine extracts as new sources for antimalarial molecules.

## 3. Experimental Section

### 3.1 Sample collection and extractions

A total of eleven specimens from eight species were collected between summer of 2010 and winter 2011 in Azores island region. Hydrothermal vent samples (mussel, shrimp and bristle worm) were collected using ROV Victor 6000 onboard R/V “Porquoi Pas?” in Lucky Strike, at about 1700 m. Specimens were immediately frozen using liquid nitrogen and kept at −80°C until being extracted. Mussels were separated in tissues – gills, digestive glands and muscle rests in a cold chamber onboard at 10°C to prevent chemical and biochemical alterations in tissues and frozen in pools using the same procedure. Sponges and corals, from depths between about 100 to 400 meters were collected accidentally (fisheries by-catch) during the annual longline surveys conducted off Azores islands onboard R/V “Arquipélago”. Samples were frozen immediately after being capture and were kept at −20°C until further analysis. Total lipids were extracted using a modified Bligh and Dyer method in lyophilized samples previously reduced to powder. The extraction methodology is described in detail elsewhere (16). Crude extracts of total lipids were obtained with quantities varying between 56 to 319 mg.

### 3.2 Biological assay: antimalarial testings

#### 3.2.1. Evaluation of antimalarial activity using WHO microtest methodology

Human malaria parasites were cultured as described by Trager and Jensen (17). The antimalarial activity of the compounds was determined according WHO microtest methodology (MARK III methodology). Briefly, stock solutions (10 mg/mL) for each extract were prepared in DMSO, and were diluted in medium to give a series of concentrations ranging from 15.625 μg/mL to 1000 μg/mL. Ninety microliters of each testing concentration, together with 10 μL of synchronized parasite cultures at ring stage with 0.6 % parasitaemia and 5% of hematocrit were distributed in duplicate, into each of the 96-well plates. Plates were incubated for 24 −30 h at 37 °C depending on the developmental stage of the trophozoites in the control well (parasite culture without extracts). After incubation, a series of thick films was prepared corresponding to each studied concentration. The thick films were stained with Giemsa stain. Post culture blood slides were examined by optical microscopy and the counts were read as the number of schizonts with three or more nuclei out of a total of 200 asexual parasites. The IC50 of individual samples for each extract was determined as the concentration that inhibited 50% of parasite development relatively to the control (free-drug). The obtained data were analyzed by nonlinear regression analysis, GraphPad Prism 5 software was used for this purpose. The IC50 values are expressed as means ± SD from two independent experiments.

#### 3.2.2. Cytotoxicity

HepG2 A16 hepatic cell line was cultured William’s E culture medium (Sigma, ref. W-4125) supplemented with 10% Fetal Bovine Serum (Sigma, ref. F7524), in a humidified atmosphere of 5% CO_2_ at 37 °C. Cell viability was determined based on MTT (Sigma, ref. M-5655) essay. Briefly, HepG2 cells were seeded in a flat-bottomed 96-well tissue culture plate at a density of 30×10^4^ cells/well and allowed to adhere overnight. Then 200 μL of medium containing seven different concentrations of each extract (concentrations ranging from 3.91 to 250 μg/mL) were added to each well, which were changed after 24h of incubation under standard culture conditions. At the end of the incubation period, MTT (thiazolyl blue, Sigma) solution (5 mg/ml) in RPMI 1640, without phenol red (Sigma, ref. R-8755) was added to each well. After incubation at 37 °C for 3 h, the supernatant was removed and 200 μl of isopropanol were measured into each well. Color development, proportional to cell growth or viability, was determined by measuring the optical density (OD) at 570 nm (ref. 630 nm) with an ELISA plate reader. Values for IC50 were determined with GraphPad Prism 5 software, as IC50 defined as the inhibitory dose that reduces the growth of the compound-exposed cells by 50%. The IC50 values are expressed as means ± SD from the three independent experiments. Selective Index was estimated to evaluate the degree of compounds selective toxicity to malaria parasites. It was calculated by the cytotoxicity IC50/ antiplasmodial IC50 ratio.

## 4. Conclusions

Marine crude extracts presented good inhibition percentages for *P*. *falciparum* strains tested (3D7 and Dd2). Due to the high spread of multi-drug resistant parasites worldwide, this results lead us to consider that more *in vivo* studies and further investigations should be address especially on parasite effect on Dd2 strain (Chloroquine resistance). The highest antimalarial activity obtained was for hydrothermal vent shrimp extract *Mirocaris fortunata*, with IC50 of 76.64 ± 6.33 μg/ml for Dd2 and 78.3 ± 1.79 μg/ml for 3D7 and deep-sea gorgonian coral *Callogorgia verticillata* with IC50 of 44.0 ± 6.1 mg/ml for Dd2 and 46.6 ± 7.64mg/ml for 3D7).

## Acknowledgments

Authors would like to thank the chief scientists and the crews at the missions ARQDAQO-32-P10, MENEZMAR, ARQDAQO-35-P11, CONDOR (1) 2011 and MOMARSAT2011, which allowed sample collection for this work and specially ROV crews from “Quest” and “Victor 6000” for their great work on samplings and photography at the hydrothermal vent sites. Also the colleagues at Azores University, who gave their precious help in the taxonomic identification for sponge and coral specimens: Raquel Pereira, Joana Xavier and Valentina de Matos. Also the curator at DOP biological collection (COLETA) Filipe Porteiro, for facilitating deep-sea materials used in this study.

The research was funded by CHEMECO (EURODEEP/0001/2007) and DEEPFUN (FCT/PTDC/MAR/111749/2009) projects and it was only possible with the cooperation of projects: CORALFISH (FP7 ENV/2007/1/213144), CONDOR (EEA Grants PT0040/2008) and ARQDAQO. This research was also supported by the funds provided by FCT/MEC to the Strategic Project (COMPETE & QREN) LARSyS Associated Laboratory & IMAR-University of the Azores.

Silvia Lino was supported by an FCT Ph.D. Grant from Fundação para a Ciência e Tecnologia (Ministry of Science and Technology of Portugal) with reference SFRH/BD/72154/2010. Ana Colaço was financed by POPH and QREN - Promoção do Emprego Científico comparticipado pelo Fundo Social Europeu e por Fundos Nacionais do MEC. S.L. and A.C. equally contributed to this work. All authors contributed to data analysis, interpretation, manuscript editing or discussions.

